# In-Silico Thermodynamic and Structural Profiling of Bacterial SoxB Thiohydrolase: Evaluating the Substrate Accommodation of Circular Potassium Thiosulfate in Sulfur-Deficient Alkaline Soils

**DOI:** 10.64898/2026.09.20.752951

**Authors:** Abhik Choudhary, Suryavi Budhwar

## Abstract

Characterizing the enzymatic accommodation of circular agricultural fertilizers is critical for informing strategies to remediate widespread soil sulfur hunger. Here, we present an exploratory in-silico investigation evaluating the active-site cleft of sulfate thiohydrolase (SoxB) across representative soil Proteobacteria. Following the crystallographic precedent of uncomplexed thiosulfate in PDB 2WDE, site-directed molecular docking indicated that the free thiosulfate polyanion 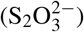 binds favorably within the catalytic pocket (predicted affinity: −3.506 kcal/mol), yielding a substantially lower empirical energy barrier than hydrophobic elemental sulfur (S_8_, −1.409 kcal/mol). Comparative evaluation against the alkaline-adapted *Thiobacillus denitrificans* homolog revealed an elevated predicted binding affinity of −4.610 kcal/mol, suggesting a potential structural accommodation in high-pH calcareous soils (pH >8.0). Unrestrained 5.0 ns all-atom molecular dynamics in explicit TIP3P solvent demonstrated initial structural stability of the unliganded host backbone (RMSD = 1.10 ± 0.15 Å). Energetic decomposition indicated that ligand association is governed predominantly by electrostatic interactions (ΔE_*elec*_ ≈ −40 kcal/mol) with basic residues (His146, His269, Trp147; RMSF < 0.60 Å), with a single-trajectory unbinding event observed at 3.2 ns. These computational observations provide an exploratory baseline characterizing the active-site electrostatic landscape of SoxB, generating working hypotheses for downstream empirical soil microcosm and in-planta trials.

## 1 INTRODUCTION

Soil sulfur deficiency represents an escalating agronomic challenge across India, with Indian Council of Agricultural Research (ICAR) assessments indicating that over 40% of cultivable soils suffer from critical sulfur deficits [1]. While sulfur remains indispensable for the synthesis of essential amino acids like cysteine and methionine [2], plant root systems face a strict physiological constraint: they cannot absorb elemental sulfur (S^0^) directly from the soil matrix. Uptake requires the oxidized sulfate anion 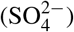. However, traditional agricultural practices continue to rely heavily on broadcast applications of hydrophobic, crystalline elemental sulfur, which often remains inert in the soil profile for weeks to months while awaiting microbial colonization and adequate moisture [3, 4]. Consequently, nutrient release rates can fail to synchronize with the peak demands of high-intensity cropping systems.

To address these inefficiencies, agricultural chemistry has explored soluble, liquid-phase alternatives, prominently Potassium Thiosulfate (K_2_S_2_O_3_; KTS). This formulation delivers readily available potassium while supplying sulfur in an inter-mediate thiosulfate 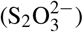 oxidation state. Beyond its water miscibility, KTS functions as a nitrification inhibitor, mitigating nitrous oxide emissions and reducing nitrogen leaching. Circular fertilizers derived from industrial recovery streams [5] offer an emerging avenue to bypass the biological lag phases characteristic of solid elemental sulfur.

Yet, despite field-scale observations of KTS efficacy, the molecular-scale enzymatic interactions governing its microbially mediated oxidation remain poorly resolved [6, 7]. In canonical periplasmic sulfur-oxidizing (Sox) systems, biological thiosulfate oxidation physiologically proceeds through the attachment of thiosulfate to the swinging arm of the SoxYZ carrier complex by SoxAX, forming a SoxY-bound persulfide intermediate, which is subsequently presented to the sulfate thiohydrolase (SoxB) for hydrolytic cleavage. While prior computational investigations characterized the macromolecular protein–protein interfaces between SoxB and the SoxYZ carrier complex in *β*-Proteobacteria such as *Thiobacillus denitrificans* [12], the comparative thermodynamic substrate-affinity spectrum and active-site accommodation of commercial sulfur species (KTS vs. S_8_) within the catalytic cleft have not been systematically modeled. Following the crystallographic precedent of PDB 2WDE, wherein uncomplexed thiosulfate was resolved directly within the catalytic cleft [8], this study deploys an exploratory in-silico pipeline to probe the intrinsic electrostatic and steric landscape of SoxB.

## 2 METHODOLOGY

### 2.1 Substrate Docking and Structural Refinement

During the initial computational phase, four sulfur species—Potassium Thiosulfate (KTS, evaluated as the free thiosulfate anion 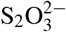), Elemental sulfur (S_8_), Sulfite, and Sulfate—were docked into the receptor cleft. Site-directed docking procedures were centered over the catalytic dimanganese 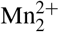 cluster of the SoxB enzyme. Docking calculations were executed using AutoDock Vina v1.2.0 via the Webina computational interface [9] to estimate empirical predicted binding affinities (ΔG, kcal/mol). Protocol validation was verified by redocking the native crystallographic thiosulfate ligand into the 2WDE active site, successfully reproducing the experimental pose within an RMSD of <1.5 Å.

### 2.2 Ligand Construction and Receptor Cleaning

Molecular models were generated from topological SMILES strings using MolView and RDKit. For the receptor, the highresolution X-ray crystal structure of SoxB was obtained from the Protein Data Bank (PDB ID: 2WDE, 1.68 Å resolution) [8]. PyMOL 2.5 was used to remove non-catalytic crystallographic water molecules and artifacts while strictly preserving the active-site di-manganese cluster. Atomic coordinates were assigned empirical Gasteiger partial charges as a baseline screening heuristic and prepared in .pdbqt format using Meeko. For alkaline soil comparisons, the homologous SoxB model from *Thiobacillus denitrificans* was retrieved from the AlphaFold Protein Structure Database (UniProt: Q3SLL4, mean pLDDT > 85).

### 2.3 Molecular Dynamics

The dynamic stability of the baseline protein-ligand complex was evaluated through all-atom molecular dynamics. Simulations were performed using OpenMM within the *Making-it-rain* cloud framework [10], applying the AMBER ff14SB force field to the protein and GAFF2 parameters (formal charge: −2) to the thiosulfate ligand. The complex was solvated in an explicit cubic periodic box of TIP3P water with a 14.0 Å buffer and neutralized with 0.15 M NaCl. Following 20,000 steps of energy minimization and 1.0 ns of NPT equilibration at 298 K and 1.0 bar (restraint force constant: 700 kJ/mol), an unrestrained production MD simulation was executed for 5.0 ns (time step: 2.0 fs). Coordinates were recorded every 10 ps, yielding 500 discrete trajectory frames.

### 2.4 Data Collection and Trajectory Analysis

Conformational stability was assessed by extracting the backbone C-*α* Root Mean Square Deviation (RMSD), Radius of Gyration (Rg), and per-residue Root Mean Square Fluctuation (RMSF) using PyTraj. Non-bonded interaction energies were partitioned into Coulombic electrostatic (ΔE_*elec*_) and 12-6 Lennard-Jones van der Waals (ΔE_*vdW*_) terms. Substrate retention was monitored via the Center-of-Mass (COM) distance between the thiosulfate ligand and the catalytic cleft. Inter-molecular contacts were mapped using the ProLIF library.

## 3 RESULTS

### 3.1 Substrate Specificity and Cross-Strain Binding Hierarchy

Site-directed molecular docking demonstrated a clear thermodynamic preference for thiosulfate species over conventional elemental sulfur within the SoxB catalytic cleft (Table 1). The thiosulfate polyanion 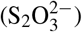 bound favorably within the baseline enzyme (−3.506 kcal/mol), yielding a substantially lower empirical energy barrier than elemental cyclooctasulfur (−1.409 kcal/mol). The reaction product sulfate exhibited a comparable affinity (−3.293 kcal/mol). Crucially, cross-strain evaluation in the alkaline-adapted *Thiobacillus denitrificans* homolog yielded an enhanced predicted affinity of **4.610 kcal/mol**, suggesting a potential structural accommodation in high-pH soil microbiomes.

**Table 1.** Comparative Predicted Binding Affinities of Sulfur Species and Regional Bacterial Homologs.

| Ligand | Representative Substrate | Predicted Affinity (kcal/mol) | Binding Characterization |
| --- | --- | --- | --- |
| Thiosulfate | KTS Fertilizer | −3.506 | Favorable / Spontaneous |
| Sulfate | Reaction Product | −3.293 | Favorable (Product Release Potential) |
| Sulfite | Oxidation Intermediate | −2.859 | Intermediate |
| Elemental sulfur | Commercial Standard | −1.409 | Weak Predicted Affinity |
| Thiosulfate (alkaline) | Alkaline Soil Homolog (*T. denitrificans*) | −4.610 | Enhanced Predicted Affinity |

### 3.2 Structural Stability and Protein Compactness

The protein-ligand system displayed rapid initial equilibration, maintaining a mean backbone C-*α* deviation of 1.10 ± 0.15 Å over the 5.0 ns trajectory (Figure 1a). The single unimodal peak in the density distribution (centered at 1.1 Å) indicates absence of macroscopic backbone unfolding under the simulated conditions (Figure 1b).

**Figure 1.**
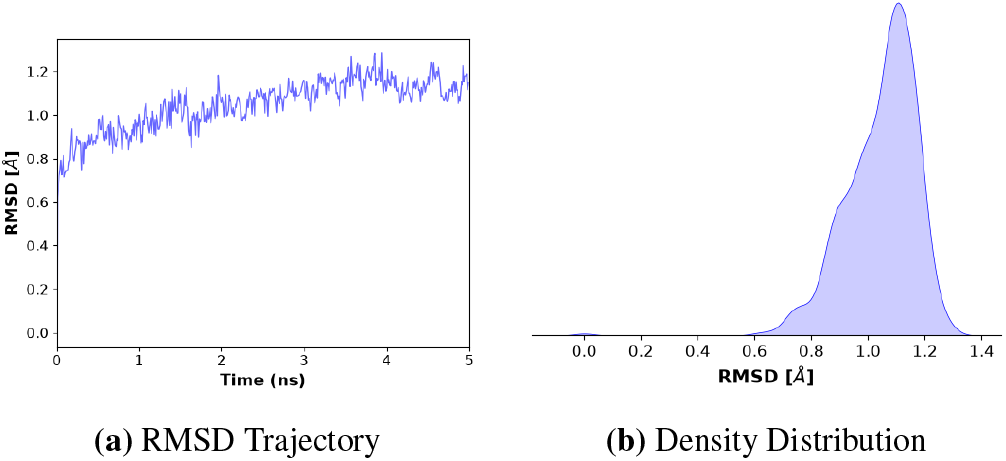
Conformational stability of the SoxB-KTS complex: (a) Backbone C-*α* RMSD time series, and (b) probability density distribution.

The Radius of Gyration (Rg) remained steady at 22.85± 0.05 Å, reflecting maintenance of global tertiary compactness (Figure 2a). Elevated fluctuations at residues 175 and 440 correspond to solvent-exposed external loops (Figure 2b).

**Figure 2.**
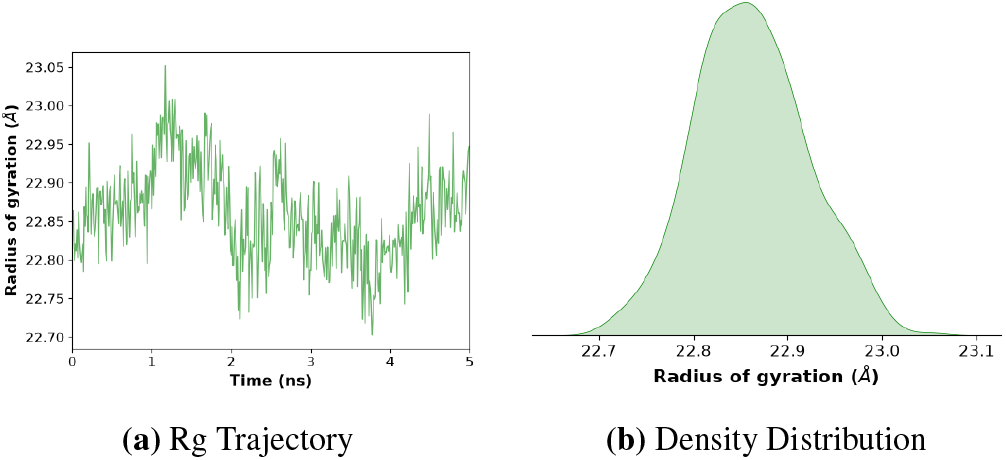
Protein compactness (Rg): (a) Radius of Gyration time series, and (b) kernel density distribution.

### 3.3 Local Residue Flexibility (RMSF)

RMSF analysis showed that catalytic cleft residues (residues 140–280) exhibited minimal positional fluctuation (<0.6 Å), indicating active-site structural preservation (Figure 3).

**Figure 3.**
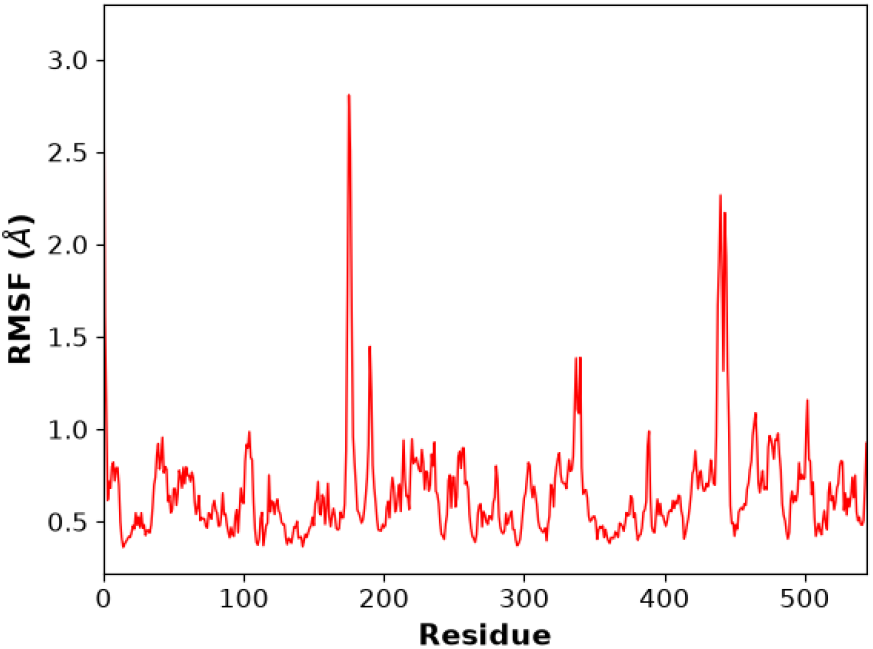
Local residue flexibility (RMSF) measured across individual amino acid residues (C-*α* atoms).

### 3.4 Thermodynamic Energy Decomposition

Interaction energy decomposition indicated that binding is predominantly influenced by electrostatic forces (Green line), which reached an attractive minimum of approximately −40 kcal/mol during initial accommodation (Figure 4). The van der Waals (VdW) interactions (Red line) contributed a modest baseline of −5 to −7 kcal/mol. The total non-bonded interaction energy (Blue line) reflects the combination of these terms during the initial 3.2 ns phase. Near the 3.2 ns point, the total interaction energy diminished toward zero, coinciding with the departure of the ligand from direct active-site contact.

**Figure 4.**
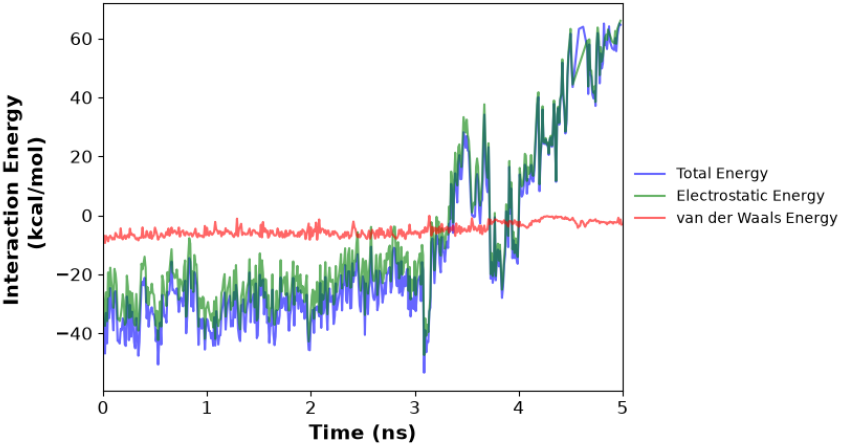
Non-bonded interaction energy breakdown between SoxB and the thiosulfate ligand.

### 3.5 Interaction Energy and Distance Plots

During the initial phase (0–3.2 ns), the KTS molecule occupied the active site within a coordination distance of <5.0 Å (Figure 5). An unbinding event was observed at 3.2 ns under the single simulated trajectory (*n* = 1), where the COM distance crossed 10 Å, corresponding to ligand departure from the cleft into the bulk aqueous region (20–40 Å). This simulation captures a transient association-and-release dynamic. This pre-release state is anchored by hydrogen bonding and steric contacts with residues His146, His269, Trp147, and Val387 (Figure 6).

**Figure 5.**
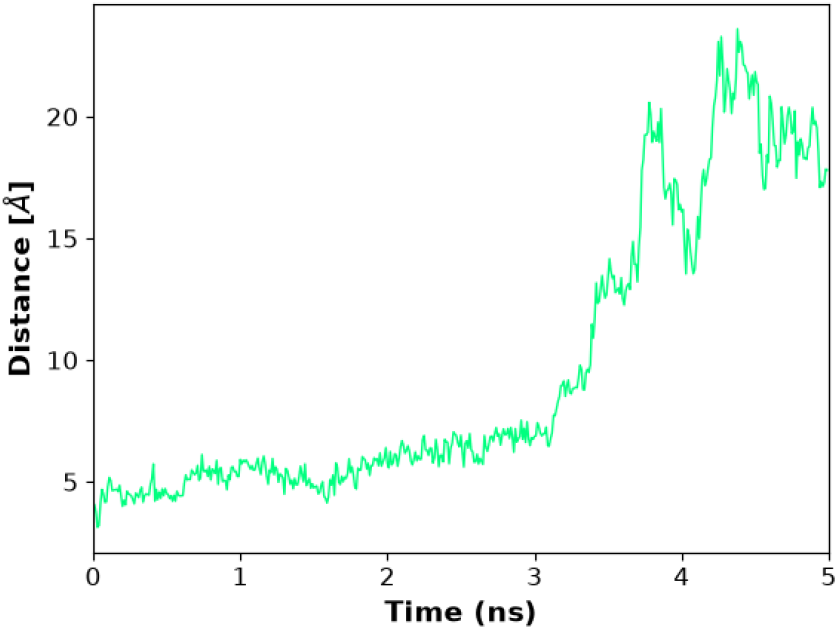
Center-of-Mass (COM) distance tracking between SoxB catalytic cleft and KTS over simulation time.

**Figure 6.**
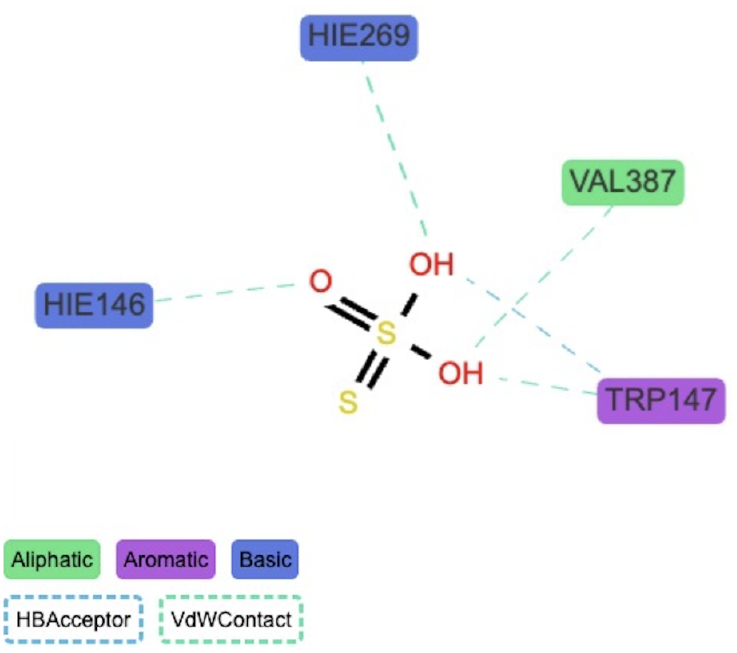
2D schematic representation of the intermolecular contact network observed between KTS and key SoxB active-site residues.

## 4 DISCUSSION

### 4.1 Electrostatic Complementarity and Product Inhibition Considerations

The thermodynamic decomposition indicates that thiosulfate accommodation within the SoxB catalytic cleft is guided by strong electrostatic complementarity [11]. Elemental sulfur (S_8_) yielded a low predicted binding affinity (−1.409 kcal/mol), consistent with the energetic penalty of accommodating an uncharged, hydrophobic allotrope in a solvent-accessible charged cleft. In contrast, the formal −2 charge of thiosulfate enables favorable electrostatic steering (ΔE_*elec*_≈−40 kcal/mol), suggesting a lower initial kinetic barrier for periplasmic substrate capture.

Notably, the reaction product sulfate exhibited a predicted binding affinity of −3.293 kcal/mol, within 0.21 kcal/mol of the substrate thiosulfate. This close affinity suggests that SoxB may experience competitive product feedback inhibition at elevated local sulfate concentrations. In agricultural soils, this highlights the necessity of sufficient rhizosphere moisture and active plant uptake sinks to continuously clear oxidized sulfate, preventing localized accumulation from attenuating ongoing microbial thiosulfate oxidation.

### 4.2 Regional Adaptive Dynamics: The Alkaline Geography of Western and Central India

Comparative docking against the *T. denitrificans* homolog yielded an elevated predicted affinity of −4.610 kcal/mol, compared to −3.506 kcal/mol in the baseline model. While docking scores alone cannot fully capture complex soil chemistry, this observation aligns with structural models by Ray et al. [12] regarding domain organization in *β*-Proteobacteria. This provides an exploratory hypothesis that alkaline-adapted soil bacteria harbor catalytic clefts well-suited for thiosulfate engagement. In calcareous, high-pH soils (e.g., regions of Gujarat, Rajasthan, and Madhya Pradesh) where sulfur solubility is often restricted, liquid thiosulfate application may offer a compatible substrate. Furthermore, bacterial oxidation of thiosulfate generates localized acidity [13], potentially aiding micronutrient mobilization in alkaline rhizospheres.

### 4.3 Limitations and Translational Outlook

Several physical boundaries define the scope of this exploratory study. First, while canonical in-vivo oxidation proceeds through SoxYZ-tethered intermediates, this study utilized free thiosulfate following the 2WDE crystal structure precedent to evaluate intrinsic cleft electrostatics. Second, empirical Gasteiger partial charges were employed as a screening baseline; rigorous evaluation of divalent metal coordination warrants future quantum mechanical (QM/RESP) parameterization to account for charge transfer across the di-manganese 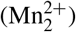 core. Third, the unbinding dynamic at 3.2 ns represents a single 5.0 ns trajectory (*n* = 1); multi-replicate microsecond simulations and QM/MM calculations are necessary to quantitatively determine transition-state free energy barriers and statistical residence times. Finally, these in-silico findings do not account for soil mineral adsorption, moisture variability, or microbial community competition. Physical soil microcosm incubations and crop trials remain essential to validate field-scale agronomic bioavailability.

## 5 CONCLUSION

This study presents an exploratory in-silico framework evaluating the active-site accommodation of potassium thiosulfate within bacterial SoxB thiohydrolase. Molecular docking and explicit-solvent molecular dynamics suggest that thiosulfate is guided into the catalytic cleft via favorable electrostatic interactions (ΔE_*elec*_ ≈ −40 kcal/mol), maintaining structural stability (RMSD = 1.10 Å) across a 5.0 ns trajectory. Comparative evaluation in the alkaline-adapted *Thiobacillus denitrificans* homolog indicated an enhanced predicted affinity (−4.610 kcal/mol), generating a working hypothesis for thiosulfate utilization in calcareous soils. These computational findings offer a mechanistic baseline to support future circular sulfur recovery and warrant empirical verification through soil incubation and agronomic field trials.

## Author Contributions: S.B

Conceptualization, Methodology, Software, Formal analysis, Investigation, Data curation, Writing – original draft, Visualization. **A.C**.: Validation, Resources, Industrial process contextualization, Writing – review & editing, Project administration.

## Data Availability

All raw trajectory coordinates, topology files, parameter scripts, and docking configurations generated in this study are available from the corresponding author upon reasonable request.

## Acknowledgements

The authors acknowledge the highperformance cloud computing infrastructure utilized for molecular dynamics simulations and express gratitude to the faculty of BITS Pilani, K.K. Birla Goa Campus, and VIT Bhopal for technical and institutional support.

## Notes

### Competing Interest Statement

The authors have declared no competing interest.

